# Polyelectrolyte complex micelles embedded in hyaluronic acid gels enable local, targeted miR-92a inhibition to accelerate diabetic wound repair

**DOI:** 10.1101/2025.11.25.690510

**Authors:** Brian Xi, Siyang Wang, Aaron Alpar, Jeffrey A. Hubbell, Matthew V. Tirrell, Yun Fang

**Affiliations:** Pritzker School of Molecular Engineering, University of Chicago, Chicago, IL 60637, USA; Department of Medicine, University of Chicago, Chicago, IL 60637, USA

**Keywords:** Actively targeted nanoparticles, polyelectrolyte complex micelles, local RNA delivery, diabetic wound healing, miR-92a

## Abstract

Diabetic wounds are characterized by various cellular deficiencies, particularly insufficient angiogenesis. MicroRNA 92a (miR-92a) is a known factor in diabetic wounds that perpetuates non-healing wound phenotypes by inhibiting angiogenesis. Therefore, its local inhibition at wound sites has therapeutic potential. To achieve this, we combine a nanoparticle formulation of polyelectrolyte complex micelles (PCMs) delivering miR-92a inhibitors with a hyaluronic acid (HA) gel formulation suitable for topical application to wound sites. The nanoparticles, formed by polyelectrolyte complexation of poly(ethylene glycol)-block-poly(L-lysine) with RNA cargo, are functionalized with targeting peptides against vascular cell adhesion molecule 1 (VCAM-1) to improve affinity for inflamed endothelial cells. We demonstrate effective PCM encapsulation and controlled release from gel formulations *in vitro* and *in vivo*. These PCMs are taken up *in vivo* by endothelial cells and exert functional transcriptional effects on miR-92a and its downstream targets. Furthermore, our composite PCM-gel formulation significantly accelerates wound closure in diabetic mouse models and improves angiogenesis, consistent with the known role of miR-92a inhibition in vascular regeneration. This work demonstrates a highly translatable formulation for improved wound healing, and lays the framework for modular nanoparticle-gel systems that can achieve local, cell-targeted RNA delivery.

**Highlights:** Polyelectrolyte complex micelles (PCMs) can be combined with hyaluronic acid gels.

VCAM-1 targeted PCMs released from gels are taken up by endothelial cells.

PCM-gels deliver miR-92a inhibitors to modulate downstream gene expression *in vivo*.

PCM-gels accelerate wound healing and enhance angiogenesis in diabetic mice.

## 1. Introduction

Non-healing wounds are significant burdens on those with diabetes. Approximately 20% of patients with diabetes experience chronic non-healing wounds, which can require aggressive interventions, such as wound debridement or even limb amputation^1–3^. Even closed wounds have high recurrence and reopening rates in those with diabetes^4^. As the global prevalence of diabetes continues to increase, so too will the need for novel therapeutics to treat diabetic wounds.

Current clinically available therapeutics primarily aim to prevent infection or maintain a moist environment to stimulate wound closure^5^. Though new treatments are currently under investigation, becaplermin, a topical gel containing recombinant human platelet-derived growth factor (PDGF), remains the only FDA-approved therapeutic for diabetic wound healing that directly modulates the wound’s cellular environment, promoting angiogenesis and stimulating production of ECM^6,7^. However, it is costly, and concerns have been raised over increased risk of cancer associated with its usage^8^.

To this end, much research has been focused on the development of novel biomaterials platforms for wound healing. Broadly, these biomaterials can be divided into two categories: nanomedicine-based therapies and hydrogel and bulk material-based therapies.

Nanomedicines, including biologically derived exosomes and synthetic nanoparticle formulations, have been effectively leveraged to improve wound healing^9–12^. Despite promising pre-clinical results, these therapies are limited by their relative difficulty of administration.

Nanomedicines that require syringe-based delivery (e.g. intravenous, subcutaneous) are not well suited for self-administration, and topical administration of aqueous formulations requires complete absorption and uptake of the nanoparticle solution into the wound area^13^, which may prove difficult for larger or irregular wounds. Additionally, nanoparticle storage and stability remain as barriers for clinical translation, particularly for RNA-encapsulating nanoparticles^14^.

Hydrogel and other bulk materials are also of key interest in treating diabetic wounds. These materials have been engineered to deliver various therapeutic cargos, such as cells and nucleic acids^15–19^, and can be intrinsically therapeutic via structural and biochemical cues^20,21^. For diabetic wounds in particular, hydrogels can act as scaffolds to promote moist healing environments and promote cellular regrowth and migration. However, the scalability and applicability of hydrogel therapeutics may limit clinical effectiveness. Hydrogels intrinsically require crosslinking to function; these crosslinking agents must be biocompatible and not interfere with wound healing, which can limit some of these novel formulations^22^. Even clinically approved crosslinkers have raised concerns due to their intrinsically reactive chemistry, requiring stringent purification before use^23,24^. In addition, formulations with complex chemistries or cellular components can be difficult to scale up for clinical use. Indeed, a vast majority of clinically approved hydrogel formulations for wound healing have relatively simple formulations for easy scalability^5,25^. Thus, despite their promising preclinical results, broader adoption of these biomaterial therapeutics has been limited.

Biologically, wound healing is characterized by four general phases. In order of occurrence after wounding, they are: hemostasis, inflammation, proliferation and remodeling^3,26^. Hemostasis, characterized primarily by clot formation, occurs rapidly within several hours of injury^27^.

Inflammation, which clears pathogens and dead cells, and the subsequent cellular proliferation proceeds in the days and weeks after wounding^28^. One key biological feature of the proliferation phase is wound angiogenesis. Capillary vessel sprouting and network formation is required to supply necessary nutrients to support granulation tissue deposition and re-epithelialization^29,30^. In standard non-diabetic wounds, angiogenesis begins later in the inflammatory phase and continues into the proliferative phase. Lastly, wound remodeling occurs over long time scales, improving the strength of newly healed skin. In this phase, the previously formed high-density capillary network undergoes a controlled regression, allowing for the wound site to return to a normal level of vessel density^31^. Diabetic wound healing is characterized by impaired angiogenesis, which limits granulation tissue formation and epithelial cell and fibroblast proliferation necessary for wound closure^32,33^. Novel therapeutics that modulate cellular pathways to promote wound repair are of key clinical interest, as evidenced by becaplermin’s commercial success.

One particularly notable therapeutic in clinical trials is MRG-110, a synthetic inhibitor of the microRNA-92a (miR-92a), which has been evaluated for both wound healing and ischemic heart disease^34^. microRNAs are small, endogenous RNAs that can broadly modulate transcriptional activity of many mRNA targets. miR-92a is an established pro-inflammatory and anti-angiogenic miRNA that is enriched in vascular endothelial cells^34–36^. As such, it is a suitable inhibitory target for diabetic wound healing, as it can modulate endothelial cell functionality and promote angiogenesis. miR-92a directly represses genes including integrin alpha 5 (ITGA5), a cell surface receptor that is critically involved in angiogenesis via promotion of cell migration^34,36^, and Krüppel-Like Factor 2 (KLF2), a critical transcription factor that maintains endothelial cell health^37,38^. Indeed, miR-92a expression is associated with upregulation of cell adhesion markers (such as vascular cell adhesion molecule 1, or VCAM-1) and pro-inflammatory cytokines in endothelial cells^38–41^. miR-92a is also known to be involved in inflammatory signaling in other cell types, such as macrophages, via exosomal exchange from endothelial cells^42^. The intravenous administration of miR-92a inhibitor MRG-110 in humans has shown suppression of miR-92a in blood in a dose-dependent manner^43^. For wound healing, MRG-110 has been shown to promote wound closure and angiogenesis after intradermal injection^35,44^. However, these studies have relied on direct cellular uptake of naked MRG-110 oligonucleotides without any delivery vehicle. The development of nanoparticle delivery platforms, particularly those targeted towards endothelial cells, hold promise in the controlled delivery of miR-92a inhibitors to inflamed vasculature, thereby reducing off-target uptake and improving efficacy and safety.

Prior studies from our lab have demonstrated the ability of a polyelectrolyte complex micelle (PCM) system to deliver miR-92a inhibitors to inflamed endothelial cells in atherosclerotic plaques *in vivo*^45,46^. These micelles consist of vascular cell adhesion molecule 1 (VCAM-1) targeting peptides conjugated to a polyethylene glycol (PEG)-b-poly-(L)-lysine copolymer, which spontaneously assemble with RNA oligonucleotides in water via electrostatic interactions. While the intravenous injections of these nanoparticles have been shown to successfully deliver miR-92a inhibitors to inflamed aortic endothelial cells to treat atherosclerosis, novel formulations are required to adapt this technology for local, topical delivery. Hyaluronic acid (HA) is a key component of the extracellular matrix, and has already been clinically approved for use in therapeutic contexts, particularly wound healing^47,48^. PCMs are particularly well suited for integration into HA gel systems, particularly over common lipid nanoparticle-based RNA delivery systems, due to their compositional simplicity, hydrophilicity, and ability to protect RNA cargo from degradation^49–51^. Specifically, our PCMs can deliver miR-92a inhibitors to inflamed endothelial cells with the aid of VCAM-1 targeting peptides decorated on their surface. As such, these PCM - HA gel composites (hereafter referred to as PCM-gels) have strong therapeutic and translational potential^52^ to improve wound healing.

In this study, we engineered and validated a novel PCM-gel complex delivery system for the local delivery of miR-92a inhibitors. These VCAM-1 targeted PCMs were used to inhibit the miR-92a signaling pathway in endothelial cells in diabetic wounds, thereby promoting angiogenesis. Local delivery was facilitated by incorporating PCMs into a HA gel matrix. Given HA’s widespread clinical use and established benefits for wound healing^47,48^, our work directly compares our formulation against a baseline HA control. A key technological innovation of this research is the integration of molecular targeting with local delivery, creating a unified system capable of local, cell-type selective transcriptional modulation after wound application. There is significant prior work demonstrating the use of peptide targeting moieties for nanoparticle delivery^45,53–55^, and several studies documenting using hydrogels for local delivery of RNA-encapsulating nanoparticles^16,56,57^. However, very few studies have combined both of these technologies into one unified platform^58,59^. Thus, our study establishes a simple, modular, and translatable system for the local, cell-type specific delivery of RNA cargo in the context of wound healing.

## 2. METHODS

### 2.1. Materials

Azide-terminated PEG2000-block-poly-L-lysine(30) was purchased from Alamanda Polymers. N-terminal dibenzocyclooctyne (DBCO)-conjugated VCAM-1 targeting peptide (sequence: VHPKQHR) was synthesized by GenScript. The miR-92a hairpin inhibitor (miRIDIAN hsa-mir-92a-3p inhibitor), miRNA inhibitor negative control, and Cy3-labeled miRNA inhibitor control were purchased from Horizon Discovery. Hyaluronic acid sodium salt was purchased from Thermo Fisher (Catalog 53747). RiboGreen assay kits were purchased from Thermo Fisher Scientific. miRNeasy mini extraction kits were purchased from Qiagen. Human aortic endothelial cells (HAECs) were purchased from Lonza, alongside EGM2 media and SingleQuot growth supplements. TaqMan miRNA primers and probes were purchased from Thermo Fisher. Other qPCR primers were ordered from IDT, and SYBR Green qPCR Master was purchased from Roche through Sigma Aldrich. Goat anti-mouse CD31 antibody was purchased from R&D Systems (catalog AF3628), and Alexa Fluor 488-conjugated donkey anti-goat IgG (A-11055) and DAPI were purchased from Thermo Fisher. For flow cytometry, DNAse 1 was purchased from MP Biomedical, and Liberase TL was purchased from Roche through Sigma Aldrich. Antibodies used in flow cytometry are listed in the supplemental documents.

### 2.2. Synthesis of VCAM-1 targeting PEG-polylysine

To conjugate the VCAM-1 targeting peptide to the aforementioned azide-terminated PEG-block-polylysine copolymer, we made use of strain-promoted azide-alkyne cycloaddition (SPAAC). A 1:1 molar ratio of polymer and peptide were mixed in DI water and allowed to react overnight at room temperature. Afterwards, dialysis using a 3kDa cutoff membrane was used to purify the reaction mixture. The purified VCAM-1 conjugated PEG-polylysine (VPP) was then lyophilized and stored at -20°C as a powder until use. To validate conjugation, ∼10mg of peptide-conjugated polymer was dissolved in 300μL of D2O and analyzed via 1H NMR spectroscopy.

### 2.3. Nanoparticle Formation and Gel Encapsulation

miR-92a hairpin inhibitor, control miRNA inhibitor, and Cy3-tagged control miRNA inhibitor were synthesized by Dharmacon and purchased from Horizon Discovery. Nanoparticles were formed as previously described. Briefly, a 2:1 mass:mass ratio of miRNA inhibitor and VPP or untargeted PEG-polylysine polymer was mixed via rapid pipetting in DI water. The resulting mixtures were allowed to incubate for 15 minutes to allow for complete complexation before being used for further downstream applications. To form composite gels, we first form a pre-gel by dissolving sodium hyaluronate (HA) at 2% (w/v) in PBS. This pre-gel solution was allowed to mix for at least 1 hour, after which fully formed nanoparticles were added to the gel. Additional PBS was added to form a final gel formulation containing 25μg/mL miRNA inhibitor in 1% HA. Gels were mixed at 4°C overnight and were stored for up to one additional day prior to use (unless otherwise specified).

### 2.4. Nanoparticle Characterization

Nanoparticle formation was characterized by DLS (Wyatt Möbius) and transmission electron microscopy (FEI Spirit 120kV). Nanoparticles encapsulating 1μg of miRNA were diluted to 100μL volume in DI water to be used for physical characterization.

To validate release from gels, a transwell assay system with 8μm pore size was used. Fully formulated PCM-gels were placed in the upper transwell insert, and nanoparticles were allowed to diffuse into the lower chamber containing either cells or PBS. To characterize gel-released nanoparticles, nanoparticle-containing PBS from the lower chamber was diluted by 100x in Milli-Q water before further use.

### 2.5. RNA Release

RNA release from the gel was tested using the transwell assay system described above. For a 24 well transwell configuration, 150μL of gels were loaded into the transwell insert and placed above a lower chamber containing 600μL PBS. At designated time points, the entire 600μL volume of PBS was collected and replaced with PBS. The freshly collected PBS then underwent RNA extraction by Qiagen miRNEasy kits to isolate miRNA from nanoparticle encapsulation^60^. RNA concentration both pre- and post-RNA purification was measured by Ribogreen assay.

### 2.6. qPCR and RNA silencing

Repression of miR-92a was validated *in vitro* using the aforementioned transwell insert system. Briefly, human aortic endothelial cells (HAECs) were seeded on the lower chamber of a 24 well plate and allowed to reach 80% confluency. Cells were then treated with 200ng/mL lipopolysaccharide for 3 hours to stimulate VCAM-1 surface protein production. Following LPS pretreatment, transwell inserts containing PCM-gels were placed above HAECs and allowed to incubate for 3 to 4.5 hours. Following this, gels were removed, and the cells were allowed to rest overnight to ensure nanoparticle uptake. RNA extraction was performed by Qiagen miRNEasy kit, and miR-92a and U6 snRNA were amplified via TaqMan RNA primers. qPCR was run on a Roche LightCycler 480 machine with TaqMan miR-92a and U6 probes. To quantify *in vivo* uptake of PCMs, chloroform-TRIzol extraction was used to isolate total wound RNA. Briefly, mouse wounds were excised, placed into TRIzol, and homogenized using Lysing Matrix D (Fisher Scientific). Chloroform was added to induce phase separation, after which the RNA-containing upper phase was collected and purified for miRNA or mRNA qPCR. Gene expression levels are calculated using the ΔΔ C_p_ (double delta C_p_) analysis, with expression being normalized to expression of glyceraldehyde 3-phosphate dehydrogenase (GAPDH) and ubiquitin (UBB). Primers are listed in the Supplementary Data Table 1.

### 2.7. *In vivo* Wound Healing Model

Mice were wounded according to the excisional splinted wound model described previously in literature^61,62^. For flow cytometric biodistribution experiments and for downstream gene expression experiments, eight-week-old C57BL/6 mice from Jackson Laboratories were used. Eight-week-old male db/db mice ordered from Jackson Laboratories were used for therapeutic wound closure experiments and angiogenesis experiments. Mice were shaved, disinfected, and placed under isoflurane anesthetic and given 0.1-0.2mg/kg intraperitoneal buprenorphine analgesic. Full-thickness wounds were made using a 6mm biopsy punch, and silicone splints (inner diameter 8mm, outer diameter 12mm) were glued into place around the wound edge to prevent panniculus carnosus-mediated wound contraction. Approximately 80μL of gel, which contains the equivalent of 2μg miR-92a inhibitor, was applied to each wound, and wounds were covered with Adaptic non-adhesive wound dressings and Tegaderm to keep gels in place and minimize infection risk. Mice were single caged following wounding, and additional buprenorphine was administered as necessary 12 and 24 hours after wounding. For db/db mice, gels were reapplied 2 days after initial wounding and gel application. Mice were sacrificed by CO2 inhalation at the experimental endpoints. C57BL/6 mice were sacrificed 24 hours after wounding and gel application for biodistribution and gene expression experiments. db/db mice were sacrificed 7 days after wounding for wound closure and angiogenesis analysis.

### 2.8. Wound Sectioning

After sacrifice, mouse wounds were excised and fixed overnight in 4% paraformaldehyde (PFA). After fixation, wounds were bisected along their longest axis and placed into cassettes with sponges and biopsy paper. These wounds were then embedded in paraffin, and serial sections were collected from the bisected wound axis. Analysis of wound closure was performed via hematoxylin and eosin (H&E) staining, as previously described^61^. Briefly, the distance between panniculus carnosus muscle endpoints was used to measure the total wound width, while the distance between epithelial tongue edges represented the extent of wound closure. Whole slide scans of stained wounds were analyzed using QuPath software^63^.

Additionally, immunofluorescence against CD31 was performed on paraffin-embedded tissue sections. Briefly, tissue sections were deparaffinized and underwent antigen retrieval. Tissues were incubated with 1:200 diluted primary CD31 antibody overnight, followed by incubation with 1:400 diluted secondary AF488-conjugated antibody. Slides were then counterstained with DAPI, mounted, and imaged via confocal microscopy. Whole tissue scans were analyzed via ImageJ. Analysis was performed by freehand selecting the wound area, excluding scab tissue (identified by the absence of DAPI signal). The selected area was then thresholded to highlight CD31^+^ vessels. Positive CD31 area was measured to determine angiogenesis.

### 2.9. Flow Cytometry

To process mouse wounds for sectioning, wounds were excised and cut into small pieces (<5mm^2^). Wounds were then digested in a solution of 0.4mg/mL Liberase TL, 14 kU/mL DNase 1, 10mM HEPES buffer, and 57.2μM β-mercaptoethanol in RPMI-1640 for 2 hours at 37°C. Wounds were then mechanically dissociated through a 70μm cell strainer, washed with 25mL of RPMI with 10% FBS, and centrifuged. The resulting cell pellet was then resuspended in FACS buffer (2% FBS with 1mM EDTA in PBS), counted, and 2 x 10^6^ cells per well were plated per well in a 96-well U-bottom plate. Cells were then washed and stained for 20 minutes in 50μL of Zombie Fixable live/dead stain and anti-CD16/32 to block Fc receptors. After another wash, cells were incubated in 50μL of antibody staining solution in FACS buffer for 30 minutes. Cells were then washed, fixed in 2% PFA in PBS for 20 minutes, and washed again prior to data acquisition. Flow cytometry was run on a Cytek Aurora spectral flow cytometer, and data were analyzed using FlowJo (BD Biosciences). Gating strategies are shown in Supplementary Figures S8.

### 2.10. Statistical Analysis

Statistical analysis was performed using Graphpad Prism 10.0. Comparisons between two groups were calculated via unpaired two-tailed t-tests, with a *p* value < 0.05 being considered significant. Comparisons between multiple groups were calculated using one-way ANOVA, with Dunnett’s post-hoc test to calculate statistical significance relative to appropriate control groups. Outlier analysis was performed via Grubbs’ test using an alpha value = 0.01 used as the cutoff. All figures and error bars show mean ± SD. Statistical significance is graphically denoted as follows: * *p* < 0.05, ** *p* < 0.01, *** *p* < 0.001, **** *p* < 0.0001.

## 3. RESULTS AND DISCUSSION

### 3.1. Preparation and characterization of VCAM-1 targeted PCM-gels

In this work, we aimed to develop a topical biomaterial for the local, targeted delivery of RNA therapeutics. To that end, we first designed polyelectrolyte complex micelle (PCM)-based nanoparticles to encapsulate our RNA cargo. Previous works have demonstrated PEG-polylysine as a modular, biocompatible delivery vehicle; additionally, surface functionalization with targeting moieties enhances cellular uptake by target cells of interest. We use SPAAC click chemistry to conjugate on an DBCO-tagged VCAM-1 targeting peptide (VHPKQHR) to an azide-terminated PEG2000-block-polylysine_30_. This conjugation is confirmed by purifying the polymer and performing NMR (Figure S1). Complexation of our synthesized VPP polymer with miR-92a inhibitors was analyzed via DLS and TEM, demonstrating consistent particle sizes after nanoparticle complexation (Figure 1C, 1E).

**Figure 1:**
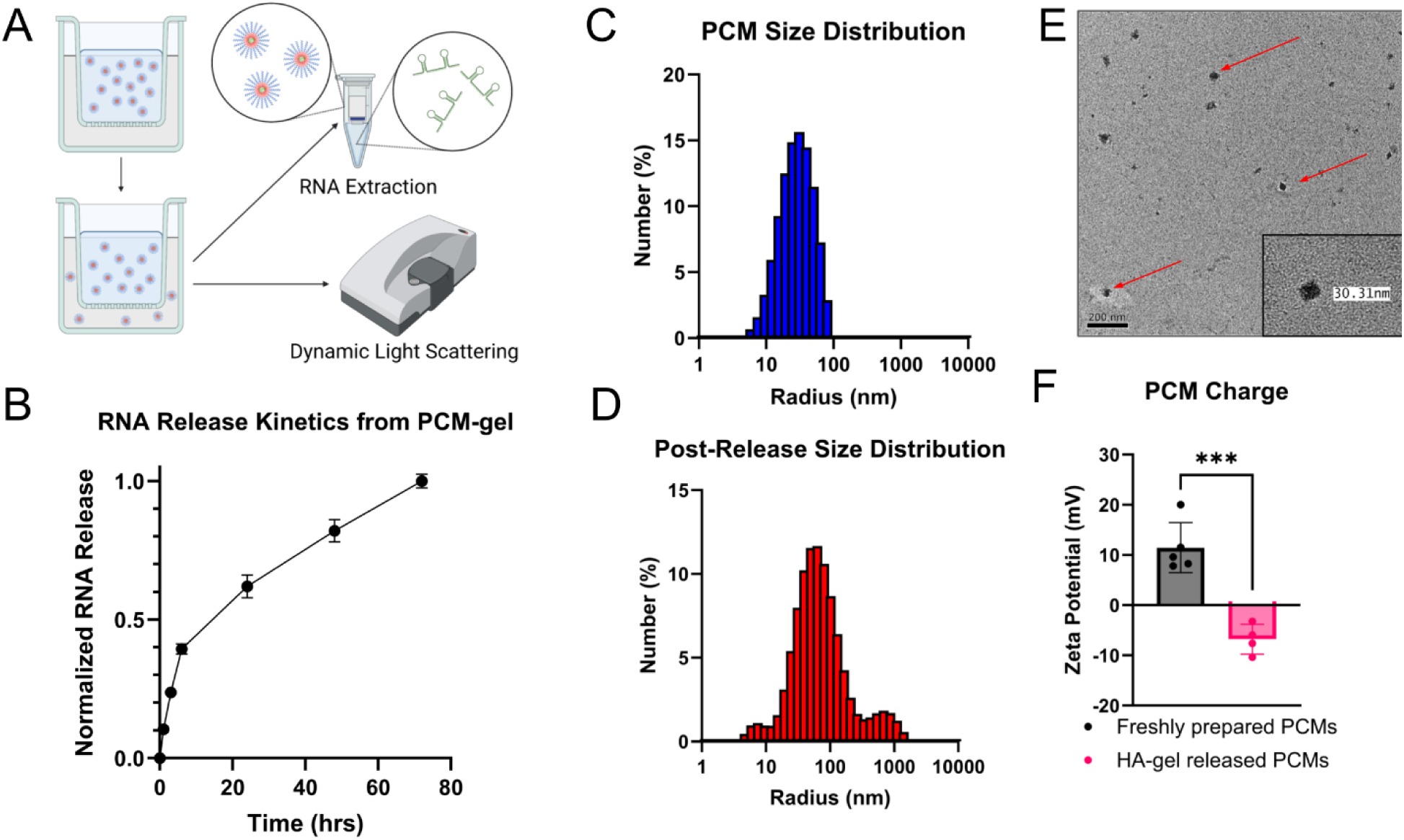
Physical characterization of polyelectrolyte complex micelles (PCMs) before and after release from hyaluronic acid gels (A) Schematic workflow for characterization of VCAM1-targeting, miRNA inhibitor-encapsulated PCMs released from HA gels via diffusion using a transwell insert. (B) miRNA inhibitor release curve shows controlled PCM release from gels over 72 hours. Data acquired by performing an RNA extraction on PCMs that have diffused from gel into PBS (n=4). (C) Size distribution of freshly prepared PCMs measured by dynamic light scattering (DLS) shows an average hydrodynamic radius of ∼30nm. (D) Size distribution of PCMs released from gel as measured by DLS shows an increased hydrodynamic radius of ∼80nm. (E) Negative-stained TEM images of freshly prepared PCMs confirm nanoparticle morphology, with sizes consistent to DLS measurements. Arrows show fully formed PCMs. (F) Zeta potentials of freshly prepared PCMs are positive, while PCMs released from gels have negative zeta potential, consistent with surface adsorption of HA to released nanoparticles (n=4-6). Data is presented as mean ± SD. Statistical analysis in (F) was performed using an unpaired Student’s *t*-test. *** *p* < 0.001

To achieve effective local nanoparticle delivery, fully complexed PCMs were mixed into a hyaluronic acid gel, yielding a final formulation containing 1% HA and a dose of 25μg/mL miRNA inhibitor. We first used a transwell membrane system to verify the release of nanoparticles from the gel into PBS, where 150 μL of PCM-gel was loaded into the upper transwell insert and the lower chamber was filled with PBS (Figure 1A). We then collected and replaced the entire PBS volume at designated time points, up to 72 hours. We then performed RNA column purification on the collected PBS to isolate the miRNA inhibitor cargo from nanoparticles and hydrolyzed HA in solution. To quantify controlled release of our RNA in this *in vitro* system we used RiboGreen assay on the purified RNA cargo (Figure 1B). This revealed a burst release followed by a more sustained release extending over more than three days.

Supplemental data shows that prior to RNA purification from the collected PBS, the amount of RNA detectable by RiboGreen assay is very low, indicating that RNA is released in an encapsulated form (Figure S2). Furthermore, DLS on released nanoparticles in PBS revealed both an increase in hydrodynamic size and decrease in zeta potential (Figure 1C, 1D, 1F); these findings indicate possible adsorption of hydrolyzed HA fragments to the surface of the nanoparticle, a phenomenon previously reported for RNA-polymer nanoparticles^64,65^.

### 3.2. RNA cargo delivery, uptake, and transcriptional effects from PCM-gels

We next evaluated whether the released nanoparticles could effectively deliver miRNA inhibitors to endothelial cells. To validate this, PCMs were formulated to encapsulate Cy3-tagged miRNA inhibitors. These nanoparticles, embedded within HA gels, were placed into the upper transwell insert and allowed to diffuse for 3 hours into the lower chamber, where human aortic endothelial cells (HAECs) had been seeded. Confocal microscopy revealed intracellular Cy3 signal in HAECs, demonstrating effective uptake of Cy3-labeled miRNA inhibitor in vascular endothelial cells from the PCM–HA gel system (Figure S3).

Additionally, we wanted to verify that our nanoparticles were indeed enriched in wound vascular endothelial cells *in vivo*. We used flow cytometry to detect and quantify the cellular uptake of our Cy3-tagged cargo in wounds. Using the splinted excisional mouse wound model^62^, we wounded 8-week-old male C57BL/6 mice and topically applied 80μL of PCM-gel containing 2μg Cy3-tagged miRNA inhibitor to their wounds immediately after the procedure. Mice were sacrificed 24 hours after gel application, and wounds were collected for single-cell isolation and downstream flow cytometry, as well as for tissue sectioning. We show a marked increase in Cy3 signal in PCM-gel treated CD31^+^ cells compared to untreated controls according to flow cytometry (Figure 2B). Additionally, we show colocalization of CD31 signal with Cy3 signal in PCM-gel treated mouse wounds via immunofluorescent staining (Figure 2C, 2D). CD31 is a marker for vascularization and angiogenesis, particularly on endothelial cells; thus, this improved uptake indicates that our nanoparticle targeting peptide can still be detected by our desired cell types. As such, these data demonstrate that PCM-gels can effectively deliver miRNA inhibitor cargo to wound vascular endothelial cells after topical administration.

**Figure 2:**
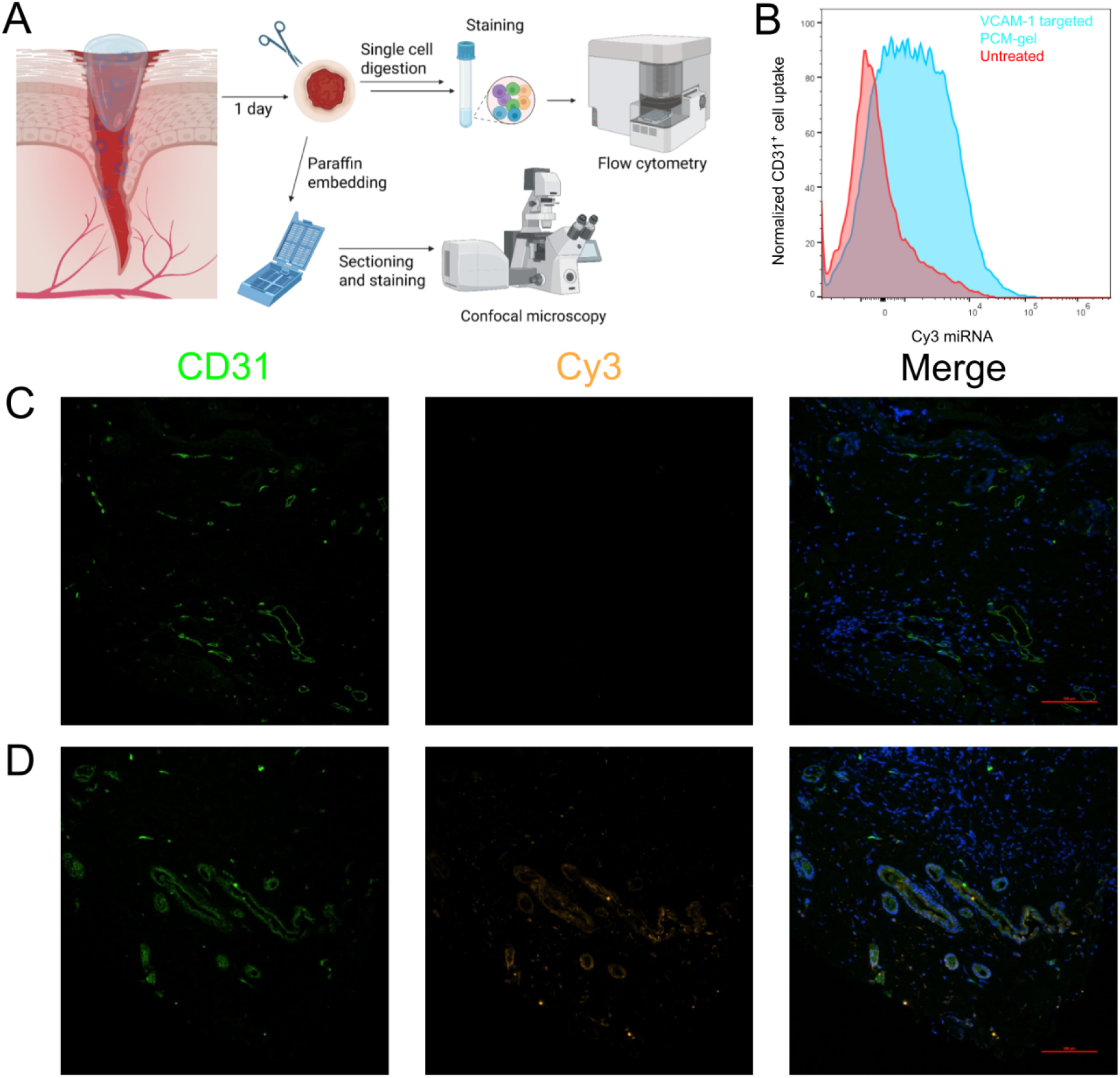
VCAM-1 targeted PCMs are enriched in endothelial cells *in vivo* after PCM-gel treatment (A) Schematic depicting biodistribution workflow after treatment of mouse wounds with PCM-gel delivering Cy3-labeled miRNA inhibitor controls. (B) Spectral flow cytometry shows that wound endothelial cells can effectively uptake VCAM-1 targeted PCMs released from gels. (C-D) Biodistribution of PCMs containing Cy3-labeled miRNA inhibitors in mouse wound sections. Green= CD31, Orange = Cy3, Blue = DAPI. Scale bar = 100µm (C) Representative section of an untreated mouse wound. (D) Representative section of a mouse wound treated with VCAM-1 targeted Cy3 PCM-gel.

Having verified that our nanoparticle-gel system was able to encapsulate and deliver intact nanoparticles, we then wanted to determine the transcriptional effects of miRNA inhibitor delivery from this system. First, we compared the miR-92a knockdown efficacy of PCM-gels with functional miR-92a inhibitors to gels containing naked, unencapsulated miR-92a inhibitors. We found that after 3 hours of gel treatment followed by overnight incubation, PCM-gels had improved knockdown of miR-92a when compared to gels containing RNA without a delivery vehicle (Figure 3B). Additionally, we tested this system *in vitro*, once again using transwell assay system to deliver functional miR-92a inhibitors or nonfunctional control miRNA inhibitors to HAECs. Our results demonstrate that PCM-gels with functional cargo can significantly inhibit miR-92a expression over control PCM-gels when treated with 150μL of gel volume (Figure 3C). Together, these data demonstrate that PCM-gels can deliver functional RNA cargo to endothelial cells *in vitro* more effectively than gels with unencapsulated RNA. Additionally, our preliminary results show that PCM-gel formulations for 25 days at -4°C are still able to retain miR-92a inhibitor functionality, supporting the translational potential of this platform (Figure S5).

**Figure 3:**
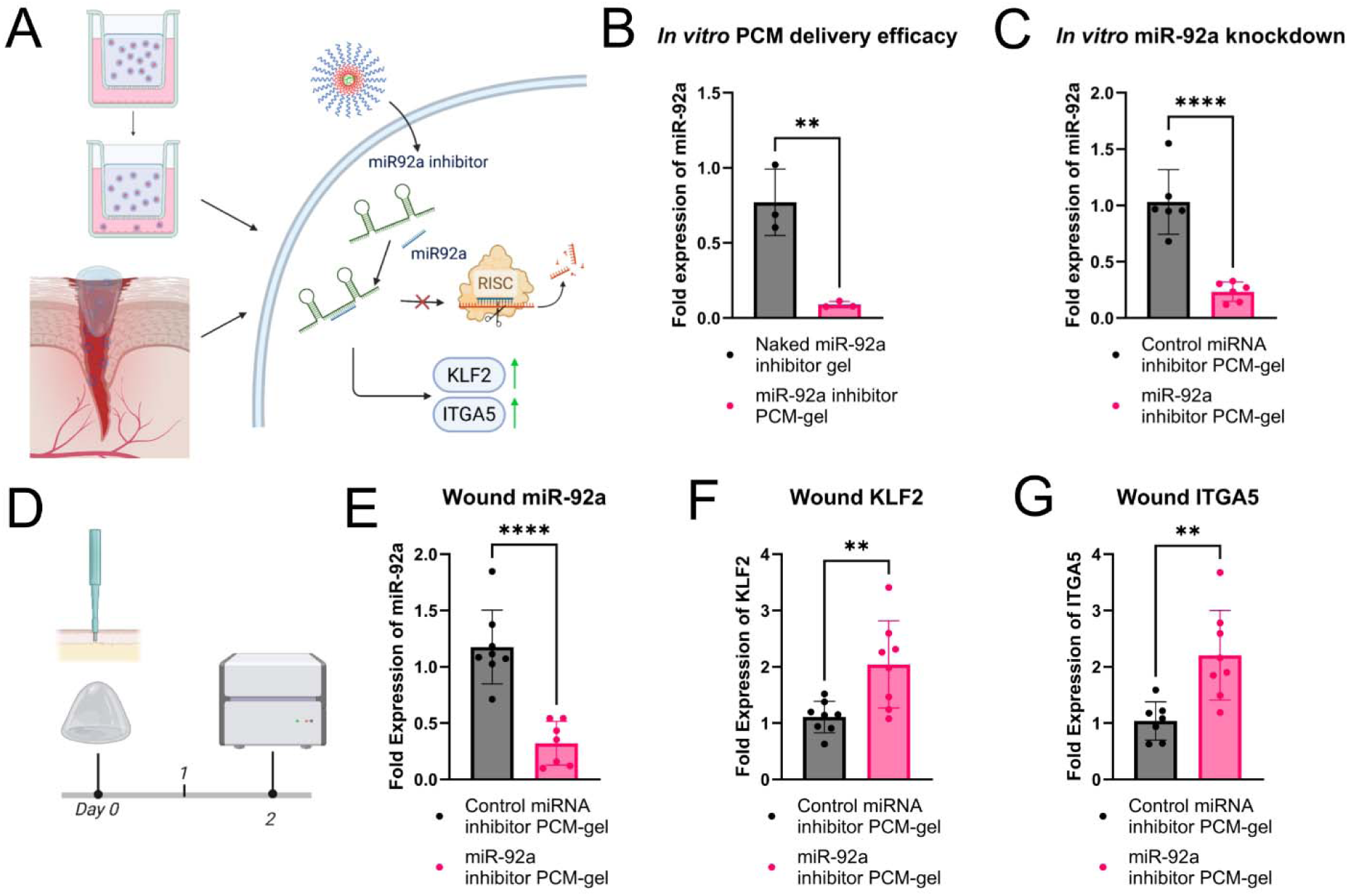
PCM-gels effectively deliver functional miR-92a inhibitor and modulate downstream gene expression. (A) Schematic illustrating the delivery of the miR-92a inhibitor by PCM-gels, tested *in vitro* using transwell cultures of endothelial cells and *in vivo* using a mouse splinted excisional wound model. (B-C) PCM-gel mediated miR-92a repression in HAECs *in vitro*. (B) PCM-gels containing VCAM-1 targeted PCMs with functional miR-92a inhibitor, loaded into the upper chamber of the transwell, significantly reduce miR-92a levels compared to gels containing unencapsulated (“naked”) miR-92a inhibitor *in vitro* (n=3) (C) PCM-gels containing VCAM-1 targeted PCMs encapsulating miR-92a inhibitor effectively decrease levels of miR-92a in HAECs compared to PCM-gels containing VCAM-1 targeted PCMs with non-functional miRNA inhibitor controls (n=6) (D-G) Delivery of functional miR-92a inhibitor by PCM-gels *in vivo* in C57BL/6 mouse wound models. (D) Timeline of mouse treatment for measuring transcriptional effects of PCM-gel applications. PCM-gels containing VCAM-1 targeted PCMs loaded with either functional miR-92a inhibitor or non-functional miRNA inhibitor controls were applied immediately after wounding. Wounds were collected 2 days after application for qPCR analysis. (E) Application of VCAM-1 targeted PCM-gels with miR-92a inhibitor to mouse wounds effectively represses miR-92a levels by ∼70% over VCAM-1 targeted PCM-gels with control inhibitors (n=7-8). (F) Wound treatment with VCAM-1 targeted PCM-gels with miR-92a inhibitor prevents the silencing of downstream transcription factor KLF2 in mouse wounds, leading to 2-fold upregulation of KLF2 mRNA levels over wounds treated with VCAM-1 targeted PCM-gels with control inhibitors (n=8) (G) VCAM-1 targeted PCM-gels with miR-92a inhibitor similarly increases ITGA5 mRNA levels compared to wounds treated with VCAM-1 targeted PCM-gels with control inhibitors (n=8). Data is presented as mean ± SD. Statistical analysis in (b), (c), (e), (f), and (g) were performed using an unpaired Student’s *t*-test. ** *p* < 0.01, **** *p* < 0.0001

*In vivo*, we demonstrate that PCM-gels can deliver miR-92a inhibitor to wounds following topical application. PCM-gels were prepared with either control miRNA inhibitor or miR-92a inhibitor, and gels were applied to mice immediately following wounding. Two days after initial gel application, we discovered that miR-92a expression in the wound is significantly repressed by miR-92a inhibitor-containing PCM-gels compared to control PCM-gels (Figure 3E). Furthermore, we found that miR-92a targets KLF2 and ITGA5 were significantly upregulated in miR-92a inhibitor PCM-gel treated wounds compared to control PCM-gel wounds according to qPCR (Figure 3F and 3G). KLF2 and ITGA5 are important factors involved in angiogenesis and wound healing and are directly downregulated by miR-92a. As such, upregulation of these genes indicates that our encapsulated miR-92a inhibitor undergoes endosomal release and has functional effects on the transcriptional level. This data supports the viability of our PCM-gel system to deliver functional miR-92a inhibitors to wound tissues and modulate relevant transcriptional pathways involved in wound healing.

### 3.3. Therapeutic evaluation of VCAM-1 targeted PCM-gels delivering miR-92a in db/db mouse excisional wound model

We then aimed to evaluate the therapeutic effectiveness of PCM-gels in promoting diabetic wound healing. As shown in Figure 4A, we surgically created full-thickness wounds in db/db mice, and PCM-gels were applied immediately after wounding and reapplied 2 days after initial application. Mice were then sacrificed seven days after initial wounding, and wounds were collected for histological analysis. We use H&E staining to detect critical biological features used to quantify wound closure (Figure 4C-D). In particular, the edges of the subcutaneous panniculus carnosus muscle are used to denote the initial width of the wounds, as they do not heal after injury; the distance between the ends of the regenerating epithelial tongue is used to determine the length of the wound that remains open. From this analysis, we show that PCM-gels with functional miR-92a cargo improve wound closure by approximately 50% over control gels with nonfunctional cargo (Figure 4B). This is particularly notable given that hyaluronic acid gels are already clinically approved for wound healing, demonstrating that PCM-gels provide significant therapeutic benefits over current treatments.

**Figure 4:**
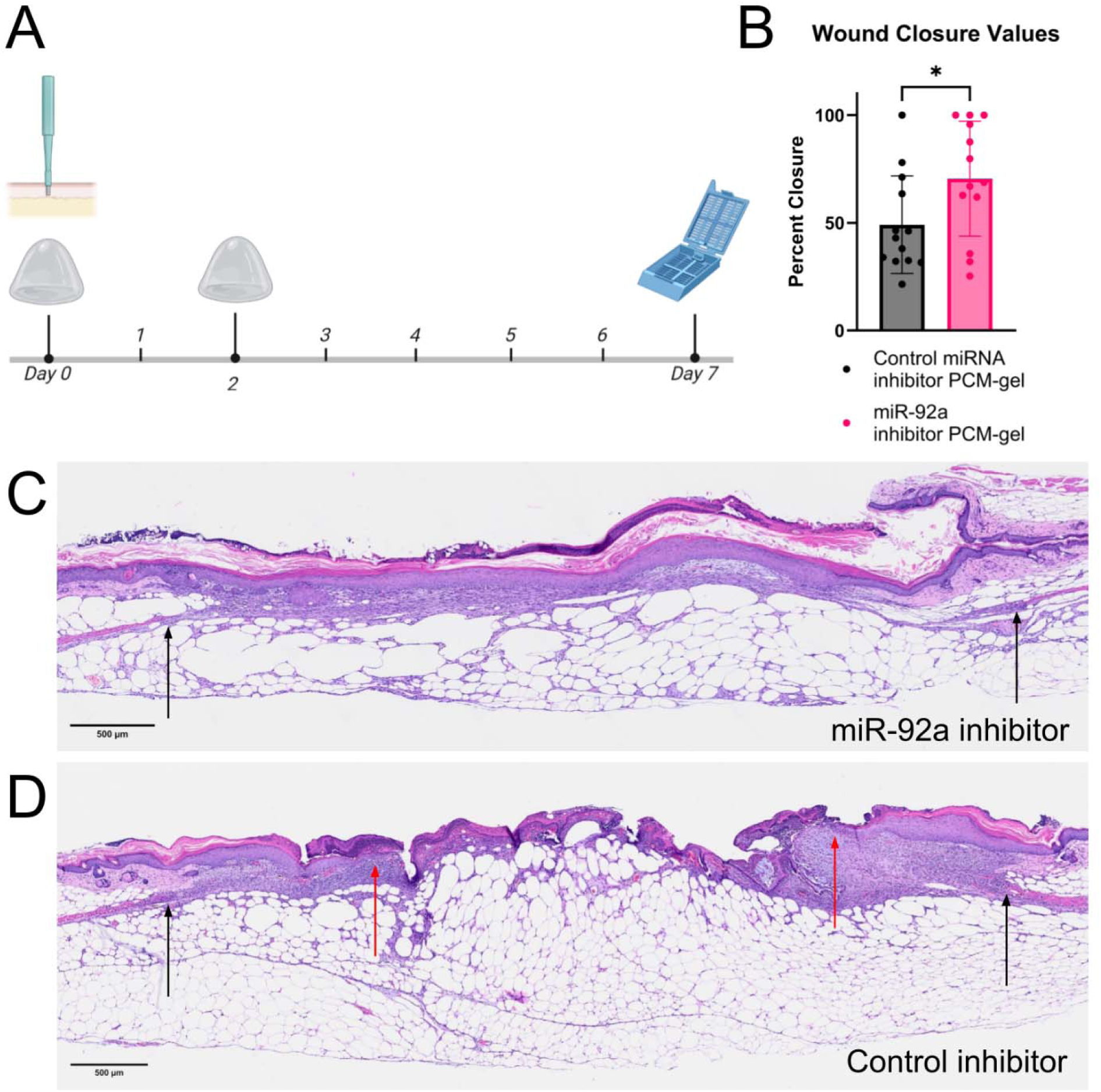
Treatment using PCM-HA gels with miR-92a inhibitor accelerate wound closure in db/db mice (A) Schematic showing PCM-gel treatment regime. VCAM-1 targeted PCM-gels containing either functional miR-92a inhibitors or non-functional control miRNA inhibitors were applied on days 0 and 2 after wounding. Mice were sacrificed for wound collection and histological processing on day 7. (B) Quantification of wound closure from H&E stain analysis shows that miR-92a inhibitor PCM-gel significantly improves total wound closure over control PCM-gels after 7 days (n=12-13 wounds). (C-D) Representative H&E sections showing wound closure of (C) <u>miR-92a inhibitor</u> PCM-gel treated wounds or (D) <u>control inhibitor</u> PCM-gel treated wounds 7 days after initial wounding. Black arrows denote original wound edges, while red arrows denote extent of reepithelialization. Data is presented as mean ± SD. Statistical analysis in (B) was performed using an unpaired Student’s *t*-test. * *p* < 0.05

To probe the cellular mechanisms for improved wound closure, we performed CD31 immunofluorescence staining at the treatment endpoint. We observed that treatment with miR-92a inhibitor-containing PCM-gels shows a significant (>50%) increase in CD31^+^ area over control inhibitor-treated wounds (Figures 5A-C). These findings are consistent with prior studies that established the anti-angiogenic activity of miR-92a. In summary, functional PCM-gels can stimulate angiogenesis via inhibition of miR-92a signaling in wounds.

**Figure 5:**
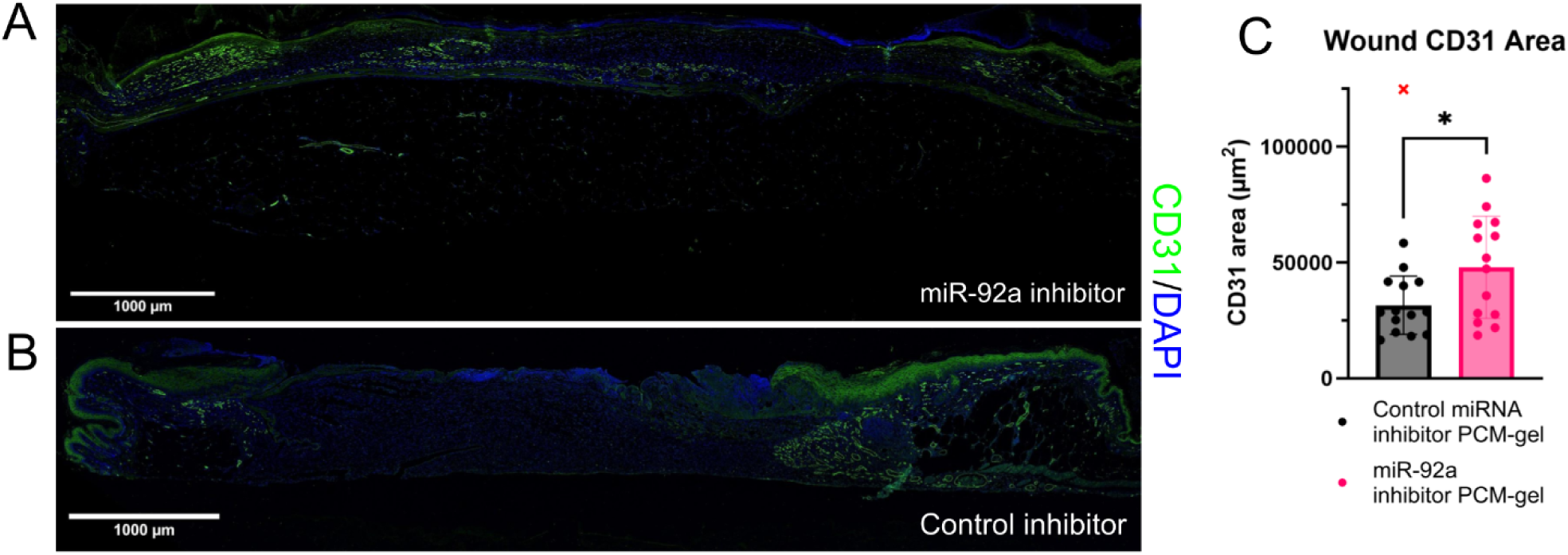
Treatment using PCM-gels with miR-92a inhibitors promotes angiogenesis in mouse wounds (A-B) CD31-stained immunofluorescent wound sections. Green = CD31, Blue = DAPI (A) Representative CD31 IF whole tissue confocal scans of <u>miR-92a inhibitor</u>-containing VCAM-1 targeted PCM-gel treated wound after 7 days of treatment. (B) Representative CD31 IF whole tissue confocal scans of <u>control inhibitor</u>-containing VCAM-1 targeted PCM-gel treated wound after 7 days of treatment. Note the minimal CD31 signal between the wound edges. (C) Quantification of CD31 area between wound edges via immunofluorescent staining shows that wounds treated with miR-92a inhibitor PCM gels have a ∼50% increase in angiogenesis compared to wounds treated with control miRNA PCM-gels. (n=14 wounds). Data is presented as mean ± SD. Statistical analyses in (C) and (E) were performed using an unpaired Student’s *t*-test. * *p* < 0.05

## 4. CONCLUSIONS

Our findings establish a three-component polyelectrolyte complex micelle (PCM)–hyaluronic acid nanoparticle-gel delivery system that can achieve local RNA delivery targeted to endothelial cells for wound repair. We engineered PCMs encapsulating miRNA inhibitor and displaying a VCAM-1 targeting peptide, which we incorporated into a hyaluronic acid gel. We then demonstrated controlled release over three days following an initial burst-release profile. Additionally, we were able to validate the encapsulation, stability, and transfection efficiency of released nanoparticles after delivery. In transwell assays, PCM-gels were able to efficiently deliver miRNA inhibitors into HAECs, thus achieving miR-92a knockdown when compared to gels delivering naked miR-92a inhibitors or nonfunctional control inhibitor-PCMs. Furthermore, these gels can be stored at 4°C for at least 25 days while still maintaining RNA functionality, supporting their translational potential. In murine wounds, we similarly found that PCM-gels achieved effective delivery to endothelial cells, reducing miR-92a levels and improving expression of miR-92a’s canonical downstream targets. Therapeutically, PCM-gels with functional miR-92a inhibitors improved wound closure by approximately 50% in db/db mice, accompanied by an approximately 50% increase in CD31⁺ area, consistent with prior findings showing the wound healing capacity of miR-92a inhibition via promotion of angiogenesis. In summary, we demonstrate that hyaluronic acid, a clinically approved material for wound healing, can be integrated with targeted PCMs to achieve a practical, translatable route for local RNA delivery for chronic wounds while overcoming issues associated with systemic nanoparticle administration.

At the core of this composite material are the polyelectrolyte complex micelle nanoparticles, which serve as targeted delivery vehicles for nucleic acid cargoes. Prior research by our lab and others have shown PCMs to be a biocompatible platform for delivery of therapeutic nucleic acids to target tissues *in vivo* following intravenous injection^45,46,66^. However, one critical limitation of these nanoparticles is their pharmacodynamic profile following intravenous administration, as the nanoparticles are cleared in a few hours post-administration. By integrating these nanoparticles into this gel formulation, we provide a means for local dosing.

The hyaluronic acid gel acts as a nanoparticle reservoir, allowing for controlled release to the target tissue of interest while minimizing opsonization and clearance in the bloodstream. Direct delivery of these nanoparticles additionally helps increase the effective concentration of therapeutics at the target site, minimizing off-target effects and improving therapeutic outcomes.

In addition to addressing pharmacokinetic limitations, the PCM–HA platform enables effective delivery of miRNA therapeutics by targeting the vascular endothelium, an essential driver of wound repair. Endothelial cells orchestrate angiogenesis, regulate inflammatory cell trafficking, and govern extracellular matrix remodeling, making them a compelling therapeutic target in chronic and diabetic wounds^33,41,67^. miR-92a is a well-characterized inhibitor of endothelial regenerative programs; its repression enhances angiogenic sprouting, improves endothelial survival, and accelerates tissue repair^34,68–70^. In support of this, layer-by-layer assembly of wound dressings delivering miR-92a inhibitors have been shown to promote wound healing, and intradermal injections of light-inducible, anti–miR-92a constructs similarly accelerate skin repair^17,71^. Together, these findings highlight the therapeutic relevance of miR-92a inhibition and underscore the importance of localized, controlled delivery of anti–miR-92a agents to effectively promote wound healing. Our strategy provides an alternative approach in which local delivery of anti–miR-92a agents is achieved through the controlled release of PCMs embedded within HA gels. By integrating endothelial-targeted nanoparticles with a clinically used HA matrix, we provide a translationally relevant strategy for restoring impaired healing in chronic wound environments.

While lipid nanoparticles (LNPs) are the current clinical standard for RNA delivery, the unique properties of PCMs make them better suited for incorporation into gel-based formulations. LNPs suffer from poor storage stability, requiring either specific buffering, ultra-low temperature storage conditions, or lyophilization to maintain efficacy over time^72–74^. These conditions render LNPs incompatible with common gel formulations, which may be incompatible with solvents and excipients required in LNP formulations, or otherwise may not tolerate LNP buffer conditions.

On the other hand, PCMs are completely water soluble and undergo self-assembly in deionized water or PBS, allowing for direct incorporation into gels without need for specialized formulations and handling. PCMs additionally have remarkable stability at a wide variety of temperatures and conditions, making them amenable for point-of-care rather than clinical use^60,75^. Overall, the ease-of-formulation of PCMs compared to LNPs indicates that PCM-gel systems may be easier to manufacture, store, and deploy than equivalent LNP-gel systems, which would aid in real-world translation of these RNA therapies.

In addition to its use in wound healing, hyaluronic acid has been clinically approved for osteoarthritis, ophthalmic procedures, and post-surgical adhesion prevention^76–78^. HA-based gels have also been widely evaluated in preclinical models of bone regeneration and cardiac repair, among other diseases^79–81^. Our results suggest that PCMs carrying RNA cargoes can be incorporated into HA to enhance their therapeutic efficacy by targeting specific transcriptional pathways involved in disease. Optimization of mechanical properties for these additional applications may require addition of non-reactive gelating agents, such as self-assembling peptides^79^.

Further work should be done to better characterize the immunological components involved in wound healing. In addition to impaired angiogenesis, diabetic wounds are characterized by persistent inflammation and impaired macrophage polarization, all of which contribute to delayed healing. In the present study, we primarily focused on the pro-angiogenic effects of miR-92a inhibition and did not directly assess how the PCM-gel influences inflammatory cell populations or macrophage phenotypes. One future direction is to modify PCMs to instead directly target immune cell signaling pathways, such as with siRNA against TNF-a, to resolve inflammatory stress. Additionally, future work must be done with regards to timing and dosing of PCM-gel applications. Given that the inflammatory phase of wound healing only starts several days after the initial wounding, our PCM-gel may be more effective if applied one to two days after the wounding^61^. However, our treatment scheme may reflect the actual application of the gel by patients, as it is likely that they treat wounds immediately rather than waiting. Further studies optimizing treatment regimen will be necessary to identify an optimal balance between efficacy and safety for these PCM-gels.

CRediT authorship contribution statement:

**Brian J. Xi:** Writing – review & editing, Writing – original draft, Visualization, Validation, Methodology, Investigation, Formal analysis, Data curation, Conceptualization

**Siyang Wang:** Methodology, Formal analysis, Investigation

**Aaron T. Alpar:** Conceptualization, Formal analysis, Methodology, Investigation

**Jeffrey A. Hubbell:** Supervision, Resources, Funding acquisition, Conceptualization. **Matthew V. Tirrell:** Writing – review & editing, Writing – original draft, Supervision, Resources, Funding acquisition, Conceptualization.

**Yun Fang**: Writing – review & editing, Writing – original draft, Supervision, Resources, Funding acquisition, Conceptualization.

## Acknowledgements

We would like to thank the support from the NIH: R35HL161244-01 (YF), HL159558-01A1 (MVT). We thank Ryne Montoya for assistance with flow cytometry. We thank the Cytometry and Antibody Technology Core Facility (Cancer Center Support Grant P30CA014599), the Human Tissue Resource Center (RRID:SCR_019199), the Integrated Light Microscopy Core (Cancer Center Support Grant 2P30CA014599), the Soft Matter Characterization Facility, and the Advanced Electron Microscopy Core (RRID:SCR_019198) at the University of Chicago. The graphical abstract and all schematic figures were made using BioRender.

---

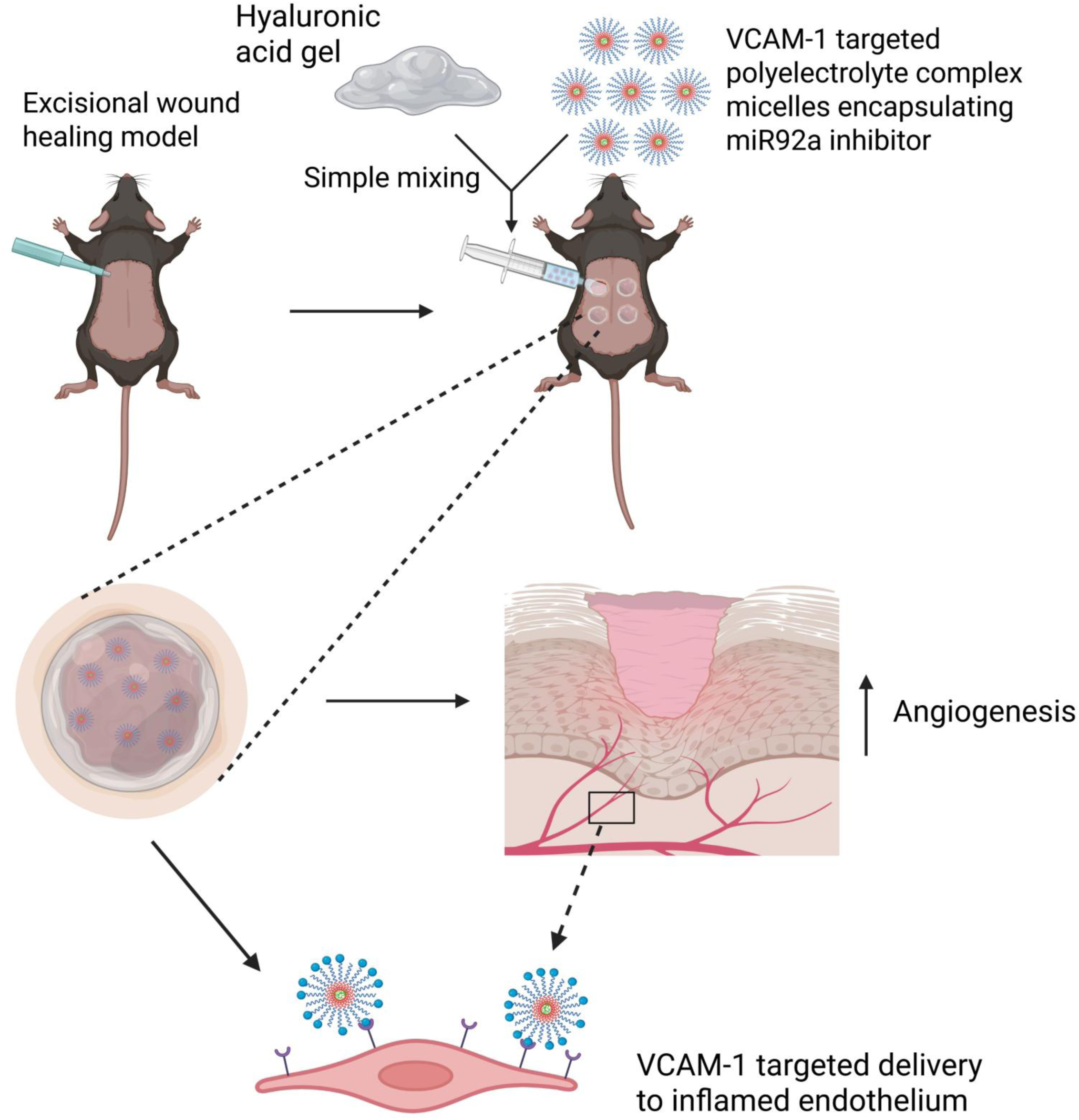

