## Supplemental Figures for "Polyelectrolyte complex micelles embedded in hyaluronic acid gels enable local, targeted miR-92a inhibition to accelerate diabetic wound repair"

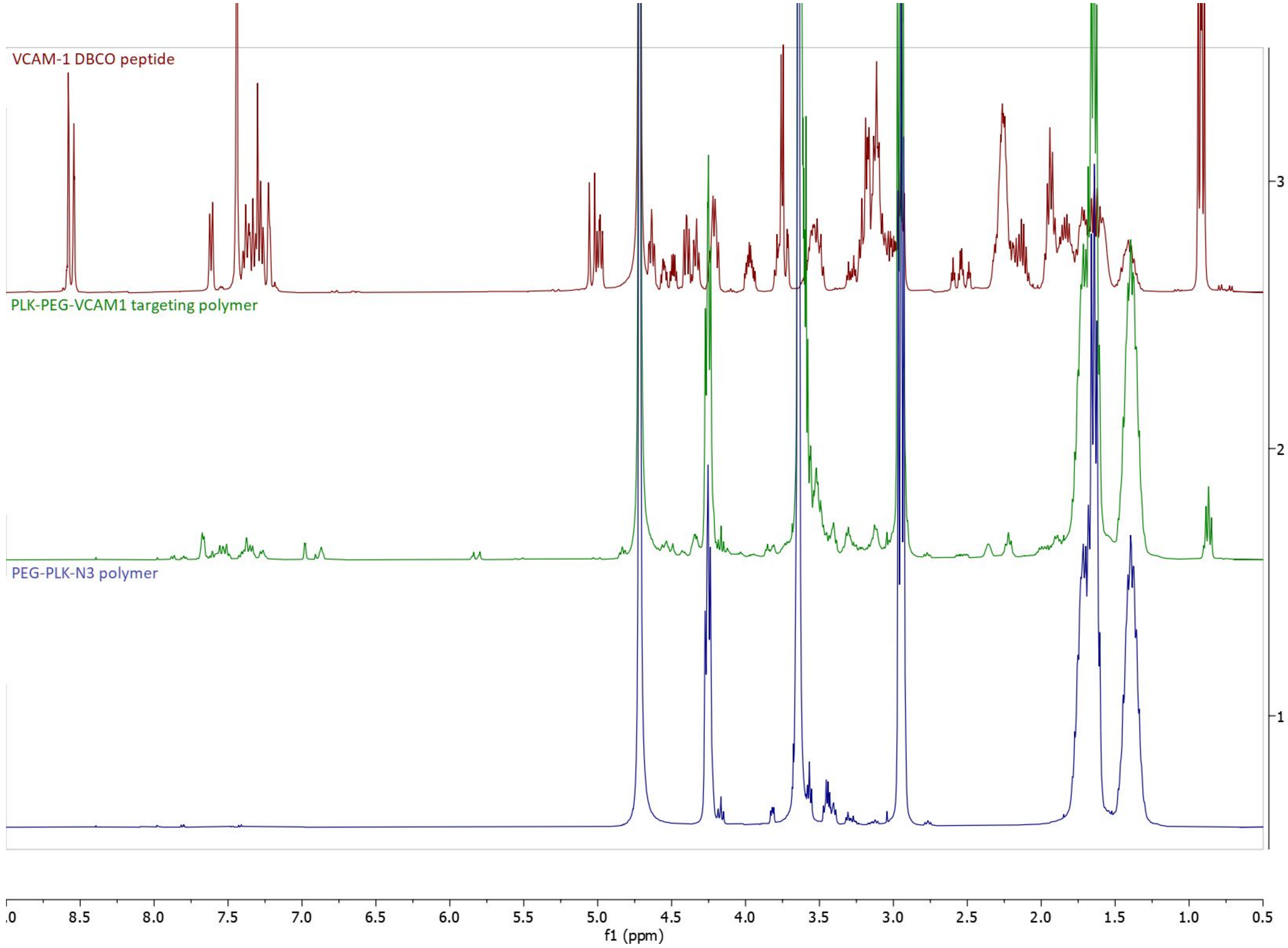


Supplementary Figure S1: NMR Showing conjugation of PEG2000-poly-L-lysine(30) with VCAM-1 targeting peptide, post purification

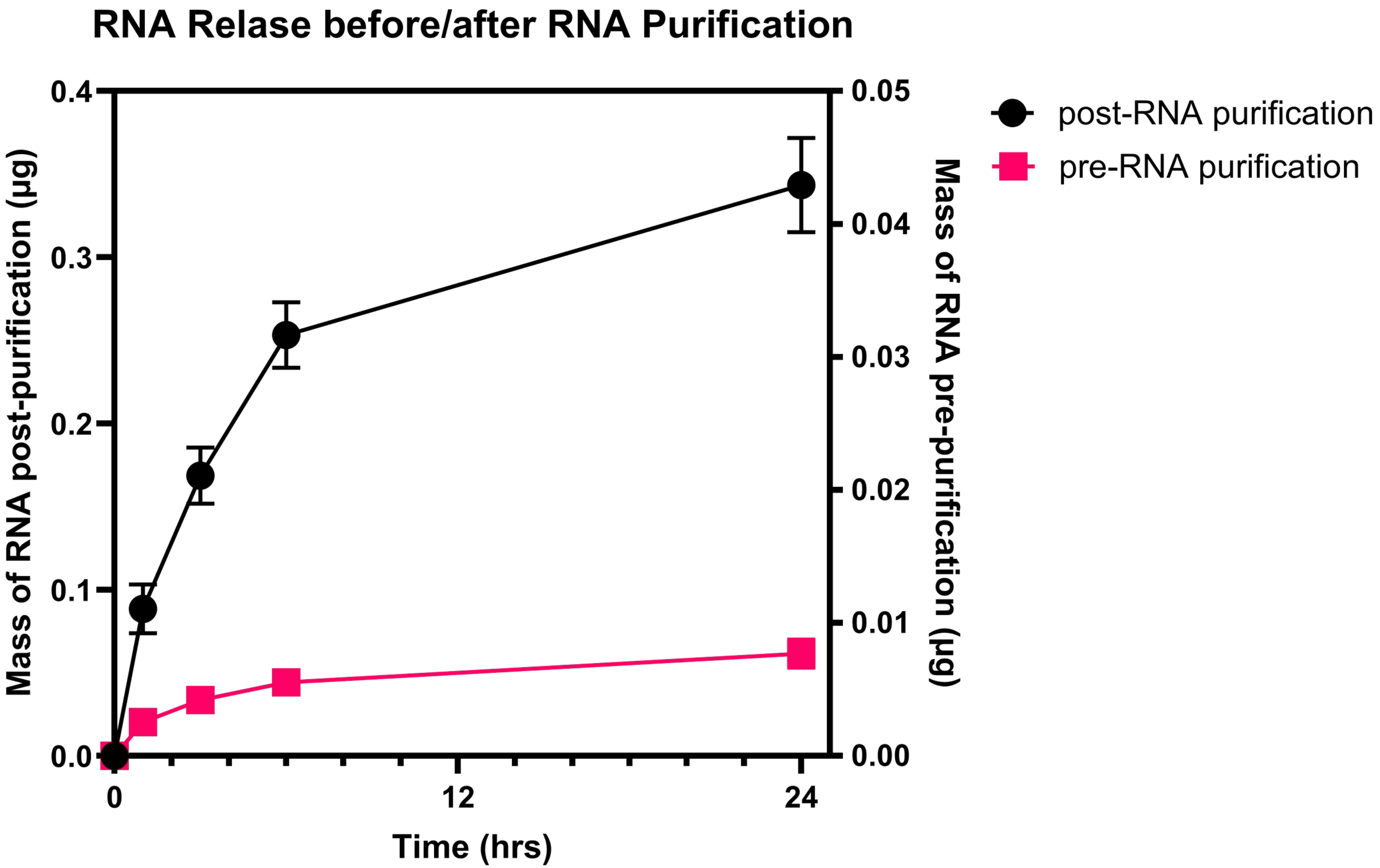


Supplementary Figure S2: RNA release curves from 150µL of gel pre- and post-RNA purification, as measured by RiboGreen Assay. Only ~2% of released RNA is detectable prior to purification, implying an encapsulation efficiency of 98% for miRNA released from PCM-gels.

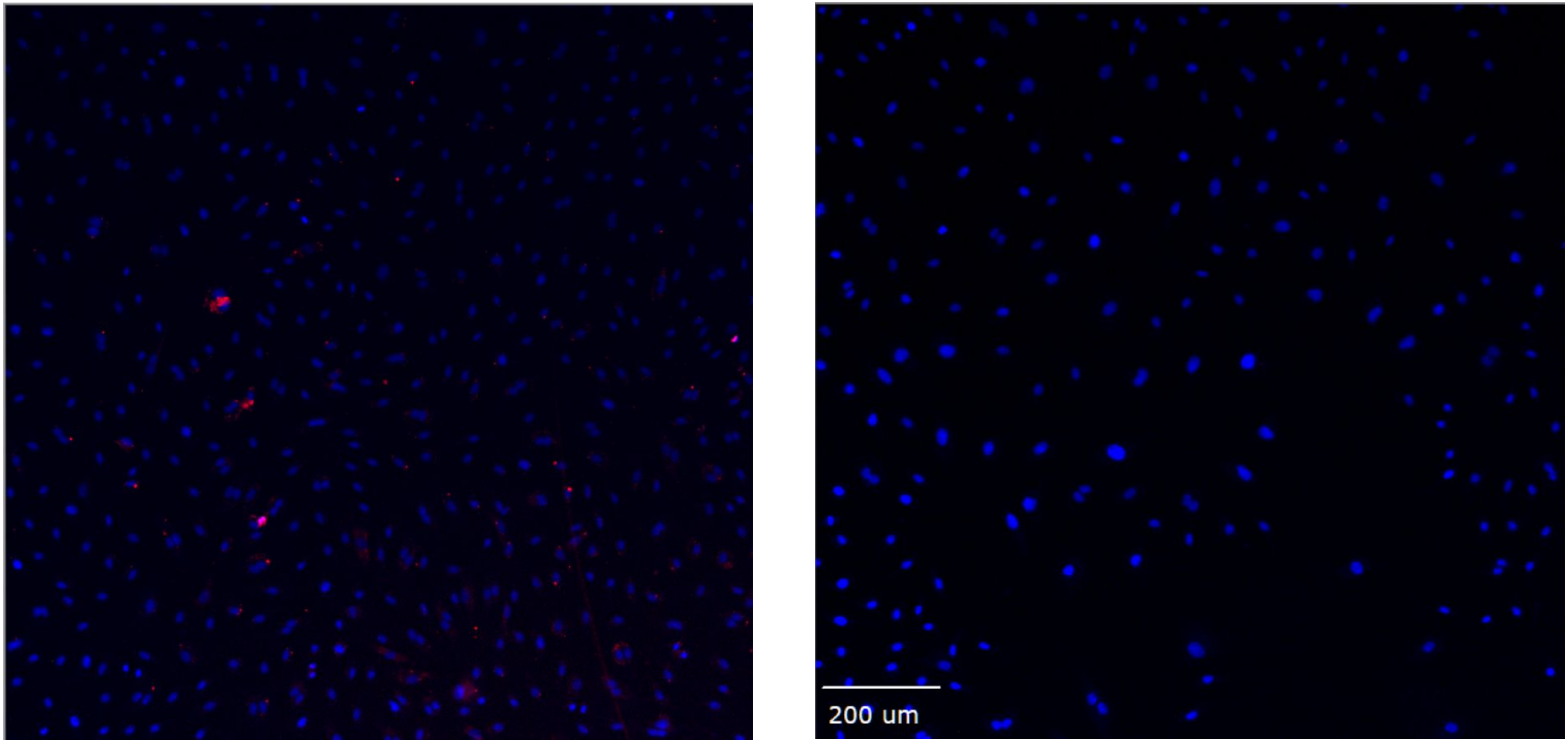


Supplementary Figure S3: Cy3-tagged miRNA inhibitor uptake in HAECs. Cells were treated for 3 hours with PCM-gels containing Cy3-labeled RNA (left) or no gels (right), and left to incubate overnight. Red = Cy3, Blue = DAPI

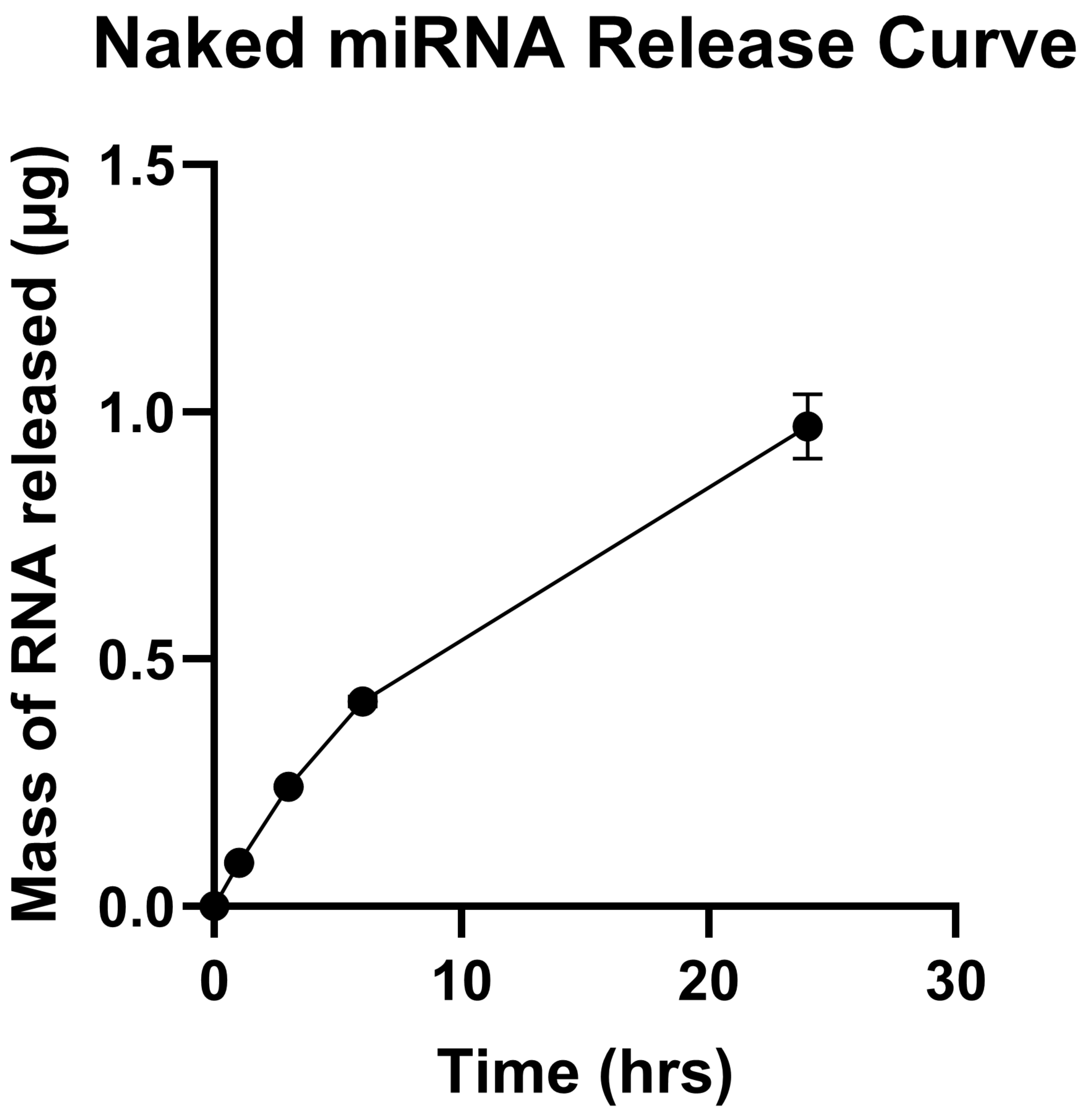


Supplementary Figure S4: Release curve for naked miRNA from gel formulation. No RNA purification was performed prior to measurement; diffused miRNA inhibitor was directly quantified by RiboGreen assay.

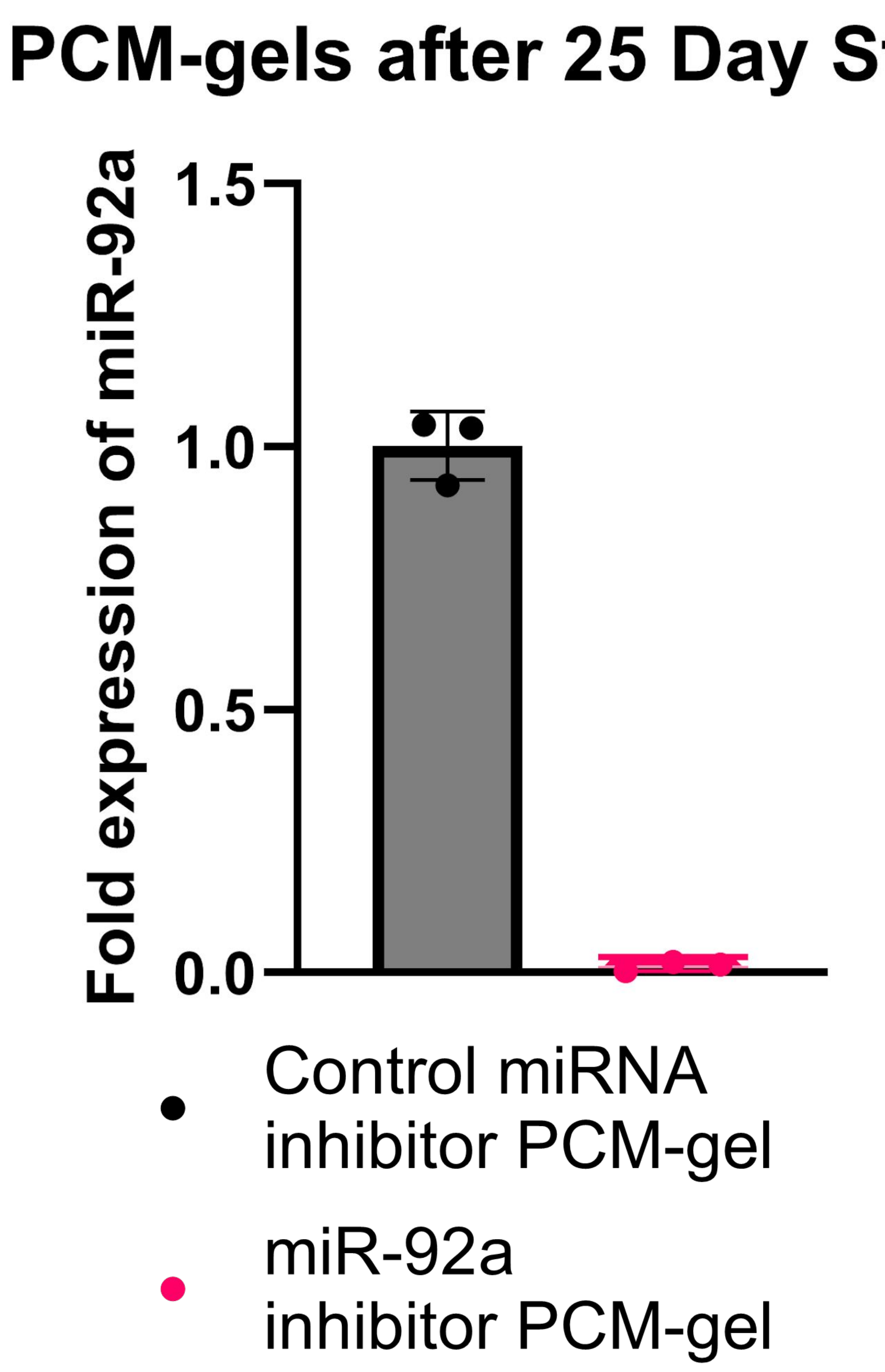


Supplementary Figure S5: miR-92a silencing efficacy of PCM-gels stored for 25 days at 4°C. HAECs were pre-treated with LPS for 3 hours, then treated with PCM-gels overnight before RNA extraction.

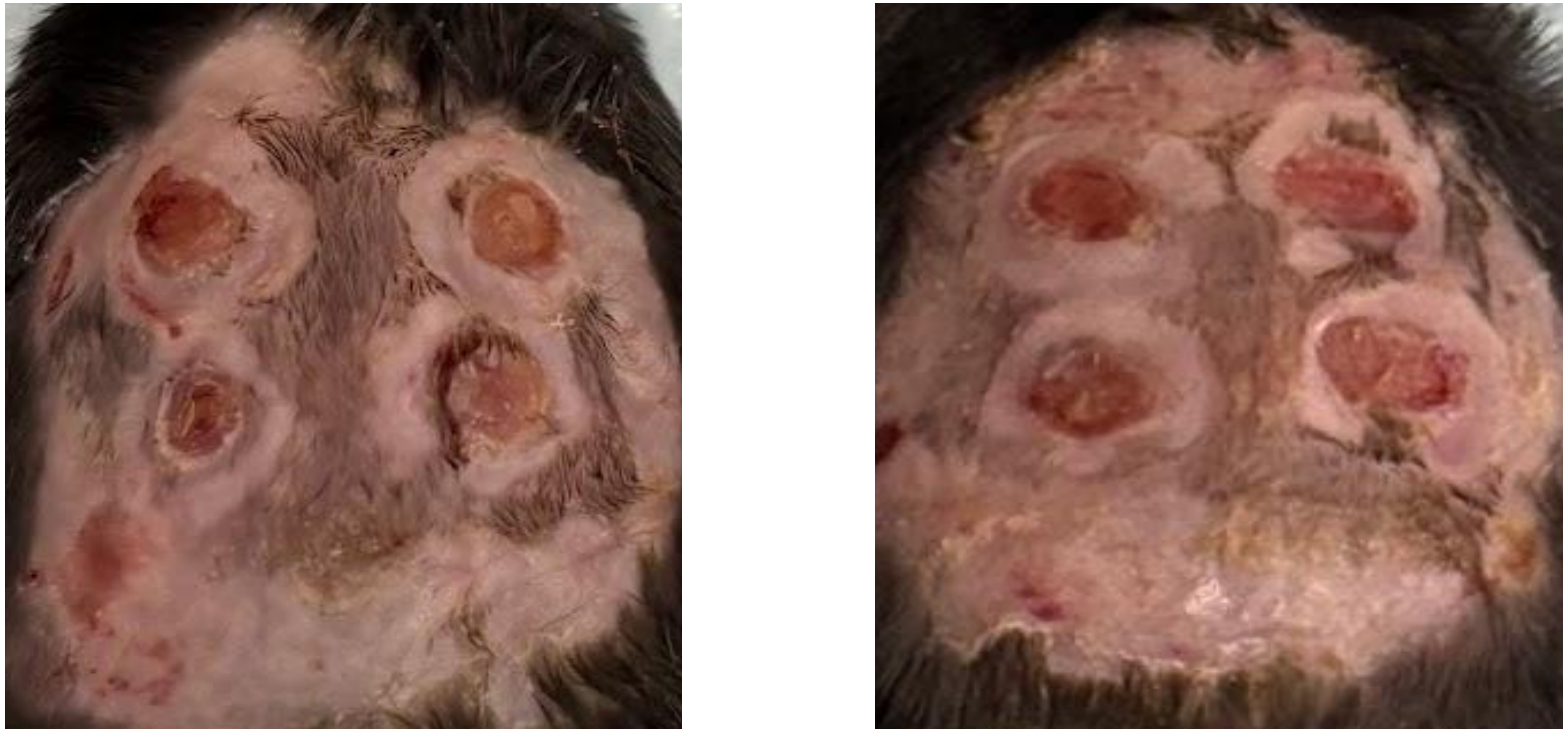

Supplementary Figure S6: Representative images of mouse wounds at time of sacrifice. Left: miR-92a inhibitor-treated mouse, Right: control inhibitor-treated mouse.

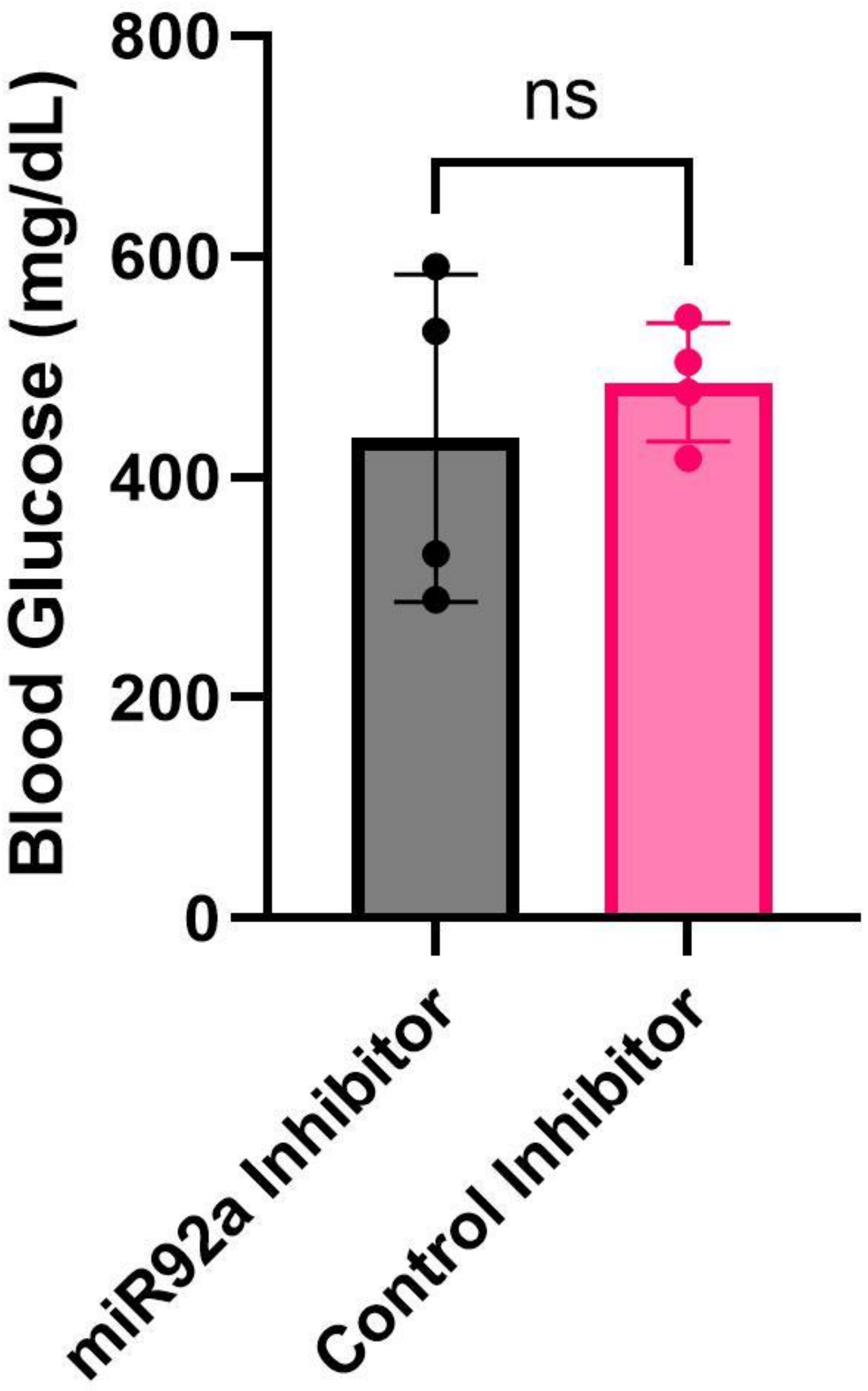


Supplementary Figure S7: Blood glucose of db/db mice used in wound closure experiments measured at sacrifice. Data from mice with blood glucose <250 mg/dL was not used.

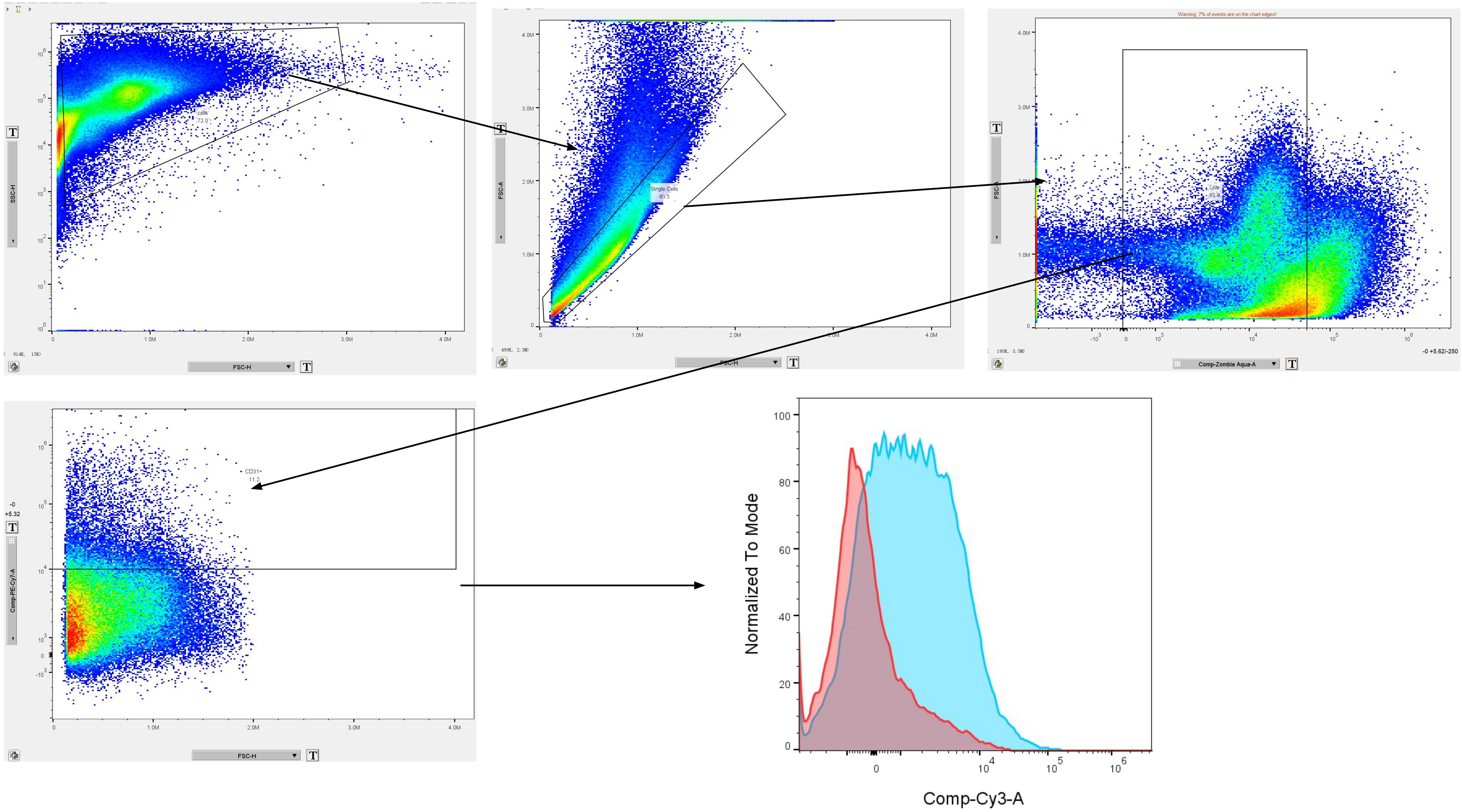


Supplementary Figure S8: Representative gating strategy for CD31 cell uptake

| Gene | Forward | Reverse |
|---|---|---|
| GAPDH | TGCACCACCAACTGCTTAGC | GGCATGGACTGTGGTCATGAG |
| UBB | AGTGACGAGAGGCTTTGTCC | CGAAGATCTGCATTTTGACCTGT |
| KLF2 | CCTTCGGTCTTTTCGAGGAC | TAAGGCTTCTCACCTGTGTGTG |
| ITGA5 | GGCACCAGTCCTATCCAGTG | GTGGAGCACATGCCAAGATG |

Supplementary Table T1: List of qPCR primers and sequences used.

| Antigen | Fluorophore | Dilution | Vendor | Catalog |
|---|---|---|---|---|
| CD16/32 | N/A | 1:200 | BioLegend | 101302 |
| Zombie | Aqua | 1:500 | BioLegend | 423101 |
| CD31 | PE-Cy7 | 1:200 | BioLegend | 102524 |
| CD45 | PerCP-Cy5.5 | 1:200 | BioLegend | 157612 |
| CD11b | FITC | 1:200 | BD | 553310 |
| F4/80 | BV421 | 1:100 | Biolegend | 123131 |
| CD206 | BV605 | 1:100 | Biolegend | 141721 |
| CD86 | BUV737 | 1:50 | BD | 741737 |
| EpCAM | PE | 1:200 | Biolegend | 118205 |
| PDGFRa | APC | 1:100 | Abcam | ab119838 |
| Ly6G | APC-Fire750 | 1:200 | Biolegend | 127651 |

Supplementary Table T2: List of flow cytometry antibodies and reagents used.
